# Missing genetic links between general factors of brain resting-state functional magnetic resonance imaging, cognition and psychopathology

**DOI:** 10.1101/2021.11.19.469227

**Authors:** JPOFT Guimaraes, B Franke, CF Beckmann, J Bralten, E Sprooten

**Author notes:** Shared Senior Authors (J Bralten, E Sprooten).

## Abstract

General factors capturing the shared genetics in psychiatric (genomic p-factor) and cognitive traits (genomic g-factor), and more recently in resting-state functional magnetic resonance imaging-derived brain networks, have contributed to our increased understanding of the etiologies in their respective domains. Yet it remains unclear whether general factors can capture the three-way genetic overlap of psychopathology, cognition and brain function. Here we tested for the presence of this genetic overlap via genetic correlation analyses using summary statistics of genome-wide association studies of the p-factor (N = 162,151 cases and 276,846 controls), the g-factor (N = 269,867), and the two genomic factors estimated from the amplitude in resting-state functional magnetic resonance imaging-derived brain networks (N = 31,688). Unlike hypothesized, only the genetic correlation between the p-factor and the g-factor was significant. We conclude that specific functional brain network constructs may have more potential than their derived general dimensions to capture relevant genetic variation for cognition and psychopathology.

## Introduction

Psychiatric disorders are highly heritable (Antilla et al., 2018). The effects of psychiatric genetic risk factors on (aberrant) behavior are associated with brain structure and functioning (Meyer-Lindenberg & Weinberger, 2006). This classic view implies that the underlying genetic mechanisms of brain function, cognition and psychopathology overlap. Yet, few studies have directly tested this three-way genetic relationship.

Recent trends in the field of psychiatric genetics have shown that the overlapping genetic influences tend to drive traits within the domain of psychopathology (Antilla et al., 2018) and the domain of cognition (Lee et al., 2012). These shared genetic influences can be reliably captured by latent factors of general genetic effects shared in each domain. This has been demonstrated for psychiatric disorders with the so-called “genomic p-factor” (Lee et al., 2019; Grotzinger, et al., 2019a), representative of the general genetic liability for psychopathology, and with the “genomic g-factor” in the case of cognitive traits (Savage et al., 2018; de la Fuente et al., 2021), which captures a shared genomic component of different higher-order cognitive functions. In a recent publication, we demonstrated that BOLD amplitudes across multiple functional brain networks, derived from resting-state functional MRI (rfMRI) scans, share genetic influences that can also be represented by latent factors (Guimaraes et al., 2021, “Shared genetic influences on resting-state functional networks of the brain,” BioRxiv, https://doi.org/10.1101/2021.02.15.431231). Specifically, two genomic factors best summarized the genetic influences shared across functional networks: one factor (F1) consisted mostly of “multimodal association” networks (ten multimodal association and two sensory networks), and the other factor (F2) consisted exclusively of “sensory” networks (six sensory networks). The overrepresentation of multimodal association networks in F1 and the exclusive association of sensory networks with F2 suggested a genetic divergence in the influence on multimodal association and sensory functions. A similar trend was also observed in a previous study using functional connectivity (FC) measures (Reineberg et al., 2020) and at the phenotypic level using network BOLD amplitudes (Bijsterbosch et al., 2017).

Literature shows that cognitive performance and psychopathology-related traits are negatively genetically correlated (Greven et al., 2011), and so are the genomic p- and g-factors (Grotzinger et al., 2019b). More recently, the p- and g-factors also were found to genetically correlate with FC between specific functional networks (Zhao et al., 2020, “Common variants contribute to intrinsic human brain functional networks,” BioRxiv, https://doi.org/10.1101/2020.07.30.229914), suggesting that the function of multiple networks shares a genetic basis with general cognition and psychopathology. Based on the previously reported intrinsic relationship between FC and BOLD amplitude (Bijsterbosch et al., 2017), and together with the evidence that different functional networks genetically overlap in their BOLD amplitudes and correlations (Elliott et al., 2018; Reineberg et al., 2020), we hypothesized that the genetic effects linking functional networks to psychopathology and cognition are shared across the BOLD amplitude of these networks and potentially represented by their general dimensions. Demonstrating this hypothesis would show that the three-way relationship between psychopathology, cognition and brain function is based on general genomic effects on brain function, in addition to effects specific to certain network constructs. Given the increased power to detect genetic overlap provided by general factors in psychopathology (Grotzinger et al., 2019a) and cognitive ability (de la Fuente et al., 2021), we expected that similar effects would exist for general factors of brain function. Here, we tested whether our two genomic functional network factors (Guimaraes et al., 2021, “Shared genetic influences on resting-state functional networks of the brain,” BioRxiv, https://doi.org/10.1101/2021.02.15.431231) and the p-(Lee et al., 2019) and g-factors (Savage et al., 2018) are genetically correlated with each other.

## Methods

### GWAS samples

To capture the genetic influences of general cognition and psychopathology, we selected previously reported GWAS summary statistics of the g-factor (N = 269,867) (Savage et al., 2018) and the p-factor coming from the latest Psychiatric Genomics Consortium cross-disorder analysis (N = 162,151 cases and 276,846 controls) aggregating eight common psychiatric disorders: anorexia nervosa, attention-deficit/hyperactivity disorder, autism spectrum disorder, bipolar disorder, major depression, obsessive-compulsive disorder, schizophrenia and Tourette’s syndrome (Lee et al., 2019). For the genetic influences shared by the activation of multiple functional brain networks, we used the GWAS summary statistics estimated for the two genomic functional network factors from our previous study (Guimaraes et al., 2021, “Shared genetic influences on resting-state functional networks of the brain,” BioRxiv, https://doi.org/10.1101/2021.02.15.431231). In brief, we derived these two factors previously using genomic structural equation modelling (genomic SEM) (Grotzinger et al., 2019a) on the summary statistics of GWAS of the BOLD amplitudes of 21 rfMRI-derived networks obtained in participants of the UK Biobank (open.win.ox.ac.uk/ukbiobank/big40/; N=31,688) (Smith et al., 2021).

### Statistical Analysis

#### Genetic correlation analysis

Taking the GWAS summary statistics of p-factor, g-factor and the two genomic functional network factors, we computed genetic correlations among these general factors using LD-score regression (LDSC) (Bulik-Sullivan et al., 2015). The bivariate LDSC calculates the genetic correlations between two traits, by comparing the SNP effect sizes reported for both traits in relation to each SNP’s linkage disequilibrium (Bulik-Sullivan et al., 2015). Significant genetic correlations were determined by Bonferroni correction for multiple comparisons (P < 0.05/6 = 0.008).

## Results

The results of the genetic correlation analyses are displayed in Table 1. The genomic p- and g-factors were negatively genetically correlated at Bonferroni-corrected significance levels (P < 0.05/6 = 0.008), validating previously reported negative genetic correlations found using different GWAS results (Grotzinger et al., 2019b). Our predominantly multimodal network factor F1 was not genetically correlated with the p- or the g-factors. Our sensory network factor F2 exhibited a nominally significant (P < 0.05) negative genetic correlation with the p-factor, but no such trend was seen with the g-factor.

**Table 1.** Correlation matrix showing the genetic correlation results obtained for the p-factor, g-factor and the two genomic functional network factors.

| | $\rho_g$ (SE) | | | | |
| --- | --- | --- | --- | --- | --- |
|  |  | F1 | F2 | g-factor | p-factor |
| $P(\rho_g)$ | F1 | - | 0.4**<br>(0.09) | -0.02<br>(0.04) | 0.05<br>(0.04) |
|  | F2 | 1.44E-05** | - | -0.02<br>(0.06) | -0.11*<br>(0.05) |
|  | g-factor | 0.62 | 0.77264 | - | -0.09**<br>(0.03) |
|  | p-factor | 0.24 | 0.029* | 0.0015** | - |

Genetic correlations of two genomic functional network factors (F1 and F2) derived from GWAS summary statistics reported by Smith et al., 2021 (Smith et al., 2021), with a genomic factor capturing general intelligence (g-factor) (Savage et al., 2018) and a genomic factor capturing general psychopathology effects (p-factor) (Lee et al., 2019). F1 comprises genetic effects shared among ten multimodal association networks and two sensory networks; and F2 is consisted of six sensory networks (for more details consult Guimaraes et al., 2021, "Shared genetic influences on resting-state functional networks of the brain," BioRxiv, https://doi.org/10.1101/2021.02.15.431231). Genetic correlations at nominal and Bonferroni-corrected ( $P < 0.05/6 = 0.008$ ) significance are respectively labeled with \* and \*\*.

## Discussion and Conclusion

The present study aimed to investigate the three-way genetic relationship of psychopathology, cognition and brain function via genetic correlations among the p-, g-factors and two factors estimated from amplitude of rfMRI-derived brain networks. Although we replicated the negative genetic correlation between the p- and g-factors, our results do not show shared genetic variation of two genomic functional network factors with either the p- or g-factors.

Our sensory network factor F2 showed a trend for a negative genetic correlation with the p-factor that was not significant after Bonferroni-correction. In our previous study (Guimaraes et al., 2021, “Shared genetic influences on resting-state functional networks of the brain,” BioRxiv, https://doi.org/10.1101/2021.02.15.431231), F2 also showed nominal (P < 0.05) genetic correlation with autism spectrum disorder (negative correlation) and with Alzheimer’s disease and body-mass index (BMI; positive correlations). Despite not being a brain-specific trait, BMI has gained relevance in the field of psychiatric genetics due to its genetic correlations with many of the disorders covered by the p-factor here (Antilla et al., 2018). An association between F2 and the p-factor would be in line with previously reported genetic correlations between the p-factor and sensory brain networks that contribute to our F2, such as the cerebellar, occipital and sensorimotor networks (Zhao et al., 2020, “Common variants contribute to intrinsic human brain functional networks,” BioRxiv, https://doi.org/10.1101/2020.07.30.229914). We speculate that some of the genetic variation driving activation in sensory networks could potentially contribute to the development of psychiatric symptomatology. Given this effect was modest and only nominally significant, follow-up studies using larger and better powered fMRI GWAS are warranted to test this hypothesis and further examine the nature of this genetic overlap between sensory network activation and psychopathology.

We did not find any genetic overlap between F1 and the p-factor, and between the g-factor and either F1 or F2. With these results, we failed to prove the three-way genetic relationship of general factors of brain function, cognition and psychopathology that we hypothesized based on previously reported genetic correlations of p- and g-factors with FC measures of individual networks (Zhao et al., 2020, “Common variants contribute to intrinsic human brain functional networks,” BioRxiv, https://doi.org/10.1101/2020.07.30.229914). Despite the intrinsic relationship between FC and BOLD amplitude (Bijsterbosch et al., 2017), our results reveal that individual network constructs such as FC perform better than our genomic factors of BOLD amplitude in capturing genetic variation relevant for the p- and g-factors. It is also possible that F1 and F2 can detect genetic overlap driven by locally relevant genetic effects (Werme et al., 2021) that may be canceled when the whole genome is analyzed.

In conclusion, our study suggests that there are no strong genetic correlations of two genomic factors of functional brain networks with general psychopathology and cognition. Even though the genomic g- and p-factors were negatively genetically correlated with each other and show better links to psychiatric disorders, we conclude that the genetic links to brain function are better captured by biologically-specific representations of the brain such as individual networks, rather than by more general representations.

## Funding

The research leading to these results received funding from the Radboud University Medical Center PhD program and the European Community’s Horizon 2020 research and innovation programme under grant agreement no. 847879 (PRIME). This work is part of the research programme *Computing Time National Computing Facilities Processing Round pilots 2018* with project reference EINF-446, which is (partly) financed by the Dutch Research Council (NWO). B.F. received additional funding from a grant for the Dutch National Science Agenda (NWA) for the NeurolabNL project (grant 400 17 602). E.S. is funded by a NARSAD Young Investigator Award (GRANT ID: 25034), a Hypatia Tenure Track Grant and Christine Mohrmann Fellowship (Radboudumc). C.F.B. gratefully acknowledges funding from the Netherlands Organisation for Scientific Research Innovation program (NWO-Vidi 864.12.003), the Wellcome Trust UK Strategic Award [098369/Z/12/Z], and the NWO Gravitation Programme Language in Interaction (grant 024.001.006). J.B. gratefully acknowledges funding from the NWO Innovation program (Veni 09150161910091).

## Acknowledgements

This work was carried out on the Dutch national e-infrastructure with the support of SURF Cooperative.

## Conflicts of interest

B.F. has received educational speaking fees from Medice. C.F.B. is director and shareholder of SBGneuro Ltd.

## Ethics approval

The UK Biobank study received ethical approval from the North West Multi-Centre Research Ethics Committee (REC reference: 16/NW/0274) and was conducted according to the principles of the Declaration of Helsinki. The cohorts included in the Psychiatric Genomics Consortium cross-disorder and the g-factor had their protocol approved by the local Ethical Committee.

## Consent to participate

‘Not applicable’

## Consent for publication

‘Not applicable’

## Availability of data and materials

The GWAS summary statistics on the amplitude of 21 rfMRI-derived brain networks are publicly available in a second release from the UK Biobank initiative via **<u>Oxford Brain Imaging Genetics Server</u>** (open.win.ox.ac.uk/ukbiobank/big40/). The GWAS summary statistics of the two genetic factors of shared genomic influences on resting-state function are made available by the corresponding author upon request. The GWAS summary statistics on the p- and g-factor are available in the Psychiatric Genomics Consortium resources (med.unc.edu/pgc/download-results/) and the Complex Genetics Lab database (ctg.cncr.nl/software/summary_statistics), respectively.

## Code availability

Software and scripts used in the genetic correlation analysis and respective quality control are made available in github.com/MichelNivard/GenomicSEM/wiki/3.-Models-without-Individual-SNP-effects.

## Author’s contributions

The conception of the idea motivating this study was elaborated by J.G., E.S. and J.B. J.G. performed the analyses, which were supervised by J.B. and E.S. J.G., E.S. and J.B. wrote the paper with contributions from the remaining coauthors.

## References

Anttila, V., Bulik-Sullivan, B., Finucane, H. K., Walters, R. K., Bras, J., Duncan, L., Escott-Price, V., Falcone, G. J., Gormley, P., Malik, R., Patsopoulos, N. A., Ripke, S., Wei, Z., Yu, D., Lee, P. H., Turley, P., Grenier-Boley, B., Chouraki, V., … Neale, B. M. (2018). Analysis of shared heritability in common disorders of the brain. Science, 360(6395). https://doi.org/10.1126/science.aap8757

Bijsterbosch, J., Harrison, S., Duff, E., Alfaro-Almagro, F., Woolrich, M., & Smith, S. (2017). Investigations into within-and between-subject resting-state amplitude variations. NeuroImage, 159, 57–69. https://doi.org/10.1016/j.neuroimage.2017.07.014

Bulik-Sullivan, B., Finucane, H. K., Anttila, V., Gusev, A., Day, F. R., Loh, P.-R., ReproGen Consortium, Psychiatric Genomics Consortium, Genetic Consortium for Anorexia Nervosa of the Wellcome Trust Case Control Consortium 3, Duncan, L., Perry, J. R. B., Patterson, N., Robinson, E. B., Daly, M. J., Price, A. L., & Neale, B. M. (2015). An atlas of genetic correlations across human diseases and traits. Nature Genetics, 47(11), 1236–1241. https://doi.org/10.1038/ng.3406

de la Fuente, J., Davies, G., Grotzinger, A. D., Tucker-Drob, E. M., & Deary, I. J. (2021). A general dimension of genetic sharing across diverse cognitive traits inferred from molecular data. Nature Human Behaviour, 5(1), 49–58. https://doi.org/10.1038/s41562-020-00936-2

Elliott, L. T., Sharp, K., Alfaro-Almagro, F., Shi, S., Miller, K. L., Douaud, G., Marchini, J., & Smith, S. M. (2018). Genome-wide association studies of brain imaging phenotypes in UK Biobank. Nature, 562(7726), 210–216. https://doi.org/10.1038/s41586-018-0571-7

Greven, C. U., Harlaar, N., Dale, P. S., & Plomin, R. (2011). Genetic Overlap between ADHD Symptoms and Reading is largely Driven by Inattentiveness rather than Hyperactivity-Impulsivity. Journal of the Canadian Academy of Child and Adolescent Psychiatry, 20(1), 6–14.

Grotzinger, A. D., Rhemtulla, M., de Vlaming, R., Ritchie, S. J., Mallard, T. T., Hill, W. D., Ip, H. F., Marioni, R. E., McIntosh, A. M., Deary, I. J., Koellinger, P. D., Harden, K. P., Nivard, M. G., & Tucker-Drob, E. M. (2019a). Genomic structural equation modelling provides insights into the multivariate genetic architecture of complex traits. Nature Human Behaviour, 3(5), 513–525. https://doi.org/10.1038/s41562-019-0566-x

Grotzinger, A. D., Cheung, A. K., Patterson, M. W., Harden, K. P., & Tucker-Drob, E. M. (2019b). Genetic and Environmental Links between: General Factors of Psychopathology and Cognitive Ability in Early Childhood. Clinical Psychological Science: A Journal of the Association for Psychological Science, 7(3), 430–444. https://doi.org/10.1177/2167702618820018

Lee, P. H., Anttila, V., Won, H., Feng, Y.-C. A., Rosenthal, J., Zhu, Z., Tucker-Drob, E. M., Nivard, M. G., Grotzinger, A. D., Posthuma, D., Wang, M. M.-J., Yu, D., Stahl, E. A., Walters, R. K., Anney, R. J. L., Duncan, L. E., Ge, T., Adolfsson, R., Banaschewski, T., … Smoller, J. W. (2019). Genomic Relationships, Novel Loci, and Pleiotropic Mechanisms across Eight Psychiatric Disorders. Cell, 179(7), 1469–1482.e11. https://doi.org/10.1016/j.cell.2019.11.020

Lee, T., Mosing, M. A., Henry, J. D., Trollor, J. N., Ames, D., Martin, N. G., Wright, M. J., Sachdev, P. S., & OATS Research Team. (2012). Genetic Influences on Four Measures of Executive Functions and Their Covariation with General Cognitive Ability: The Older Australian Twins Study. Behavior Genetics, 42(4), 528–538. https://doi.org/10.1007/s10519-012-9526-1

Meyer-Lindenberg, A., & Weinberger, D. R. (2006). Intermediate phenotypes and genetic mechanisms of psychiatric disorders. Nature Reviews Neuroscience, 7(10), 818–827. https://doi.org/10.1038/nrn1993

Reineberg, A. E., Hatoum, A. S., Hewitt, J. K., Banich, M. T., & Friedman, N. P. (2020). Genetic and Environmental Influence on the Human Functional Connectome. Cerebral Cortex, 30(4), 2099–2113. https://doi.org/10.1093/cercor/bhz225

Savage, J. E., Jansen, P. R., Stringer, S., Watanabe, K., Bryois, J., de Leeuw, C. A., Nagel, M., Awasthi, S., Barr, P. B., Coleman, J. R. I., Grasby, K. L., Hammerschlag, A. R., Kaminski, J. A., Karlsson, R., Krapohl, E., Lam, M., Nygaard, M., Reynolds, C. A., Trampush, J. W., … Posthuma, D. (2018). Genome-wide association meta-analysis in 269,867 individuals identifies new genetic and functional links to intelligence. Nature Genetics, 50(7), 912–919. https://doi.org/10.1038/s41588-018-0152-6

Smith, S. M., Douaud, G., Chen, W., Hanayik, T., Alfaro-Almagro, F., Sharp, K., & Elliott, L. T. (2021). An expanded set of genome-wide association studies of brain imaging phenotypes in UK Biobank. Nature Neuroscience, 24(5), 737–745. https://doi.org/10.1038/s41593-021-00826-4

Werme, J., Sluis, S. van der, Posthuma, D., & Leeuw, C. A. de. (2021). LAVA: An integrated framework for local genetic correlation analysis (p. 2020.12.31.424652). https://doi.org/10.1101/2020.12.31.424652

